# Role of distinct fibroblast lineages and immune cells in dermal repair following UV radiation induced tissue damage

**DOI:** 10.1101/2021.06.08.447606

**Authors:** Emanuel Rognoni, Georgina Goss, Toru Hiratsuka, Katharina I Kober, Prudence PokWai Lui, Victoria SK Tsang, Nathan J Hawkshaw, Suzanne M Pilkington, Kalle H Sipilä, Inchul Cho, Niwa Ali, Lesley E Rhodes, Fiona M Watt

**Author notes:** Corresponding author (E.R.); (F.M.W.).

## Abstract

Solar ultraviolet radiation (UVR) is a major source of skin damage, resulting in inflammation, premature ageing and cancer. While several UVR-induced changes, including extracellular matrix reorganisation and epidermal DNA damage, have been documented, the role of different fibroblast lineages and their communication with immune cells has not been explored. We show that acute and chronic UVR exposure led to selective loss of fibroblasts from the upper dermis in human and mouse skin. Lineage tracing and in vivo live imaging revealed that repair following acute UVR is predominantly mediated by papillary fibroblast proliferation and migration. In contrast, chronic UVR exposure led to a permanent loss of papillary fibroblasts, with expansion of fibroblast membrane protrusions partially compensating for the reduction in cell number. Although UVR strongly activated Wnt-signalling in skin, stimulation of fibroblast proliferation by epidermal β-catenin stabilisation did not support papillary dermis repair. Acute UVR triggered an infiltrate of neutrophils and T cell subpopulations and increased pro-inflammatory prostaglandin signalling in skin. Depletion of CD4 and CD8 positive cells resulted in increased papillary fibroblast depletion, which correlated with an increase in DNA damage and reduction in fibroblast proliferation. Conversely, topical COX-2 inhibition prevented fibroblast depletion and neutrophil infiltration after UVR. We conclude that loss of papillary fibroblasts is primarily induced by a deregulated inflammatory response, with infiltrating T cells supporting fibroblast survival upon UVR-induced environmental stress.

**Graphical abstract:** 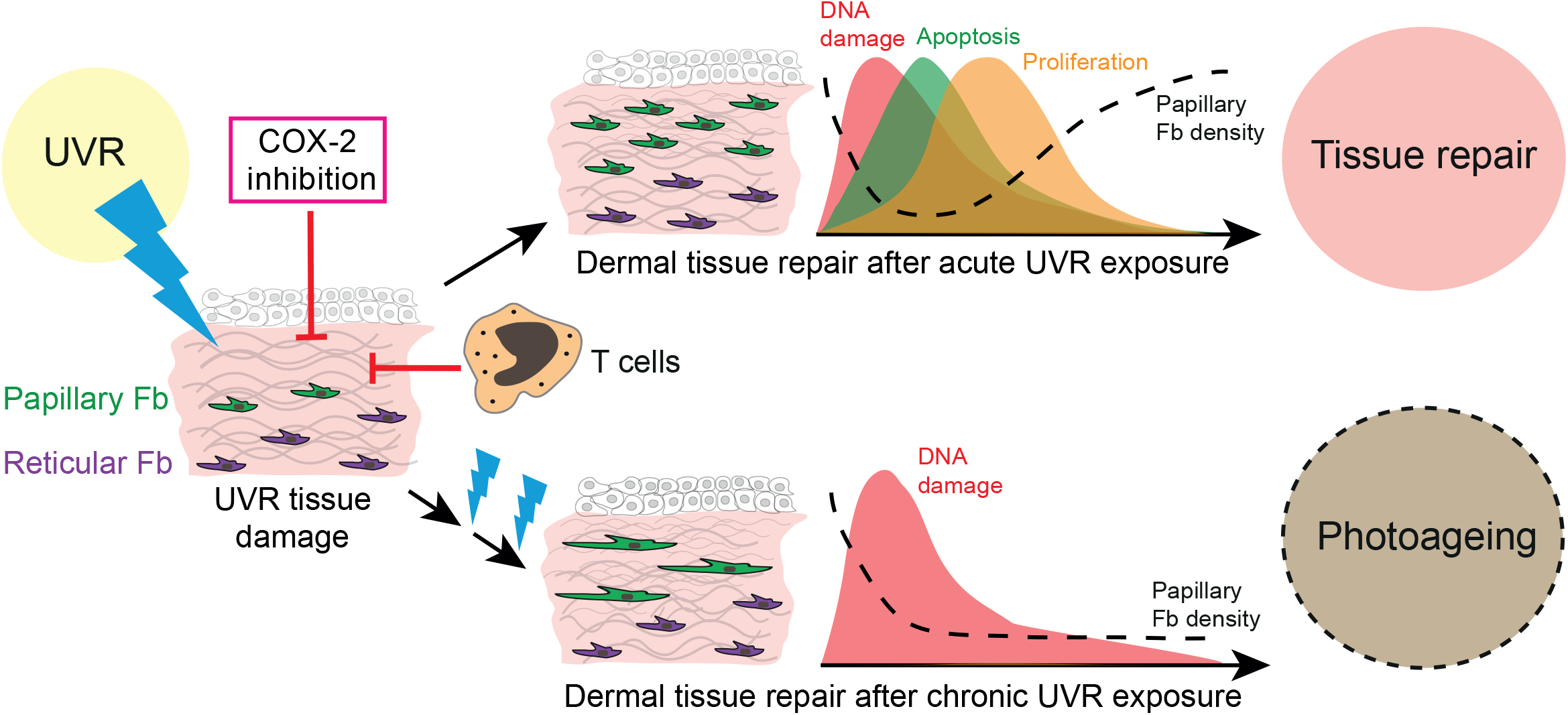

## Introduction

Ultraviolet radiation (UVR) from the sun penetrates the skin and has both positive and negative impacts on human health (1). While UVR is essential for vitamin D synthesis, prolonged (chronic) UVR exposure contributes to the development of skin cancer (photo-carcinogenesis) and accelerates ageing (photo-ageing) (2, 3). UVR is a small component of solar radiation and comprises high-energy UVC (wavelength 100–280 nm), lower-energy UVB (280–315 nm) and UVA (315–400 nm) wavebands. UVC is absorbed by stratospheric ozone while UVA and UVB penetrate the skin. Chronic UVR damages the DNA, lipids and proteins of skin cells directly (photochemical reactions) or indirectly via inflammation, reactive oxygen species (ROS) production and matrix metalloproteinase (MMP) secretion (3–6).

The connective tissue of the skin, the dermis, comprises distinct layers known as the papillary, reticular and dermal white adipose tissue (DWAT) layers (7, 8). During mouse skin development multipotent fibroblasts differentiate into distinct subpopulations (lineages) that form the different layers. These fibroblast lineages differ in location and function, and their cell identity and composition change with age (9, 10). While papillary fibroblasts beneath the basement membrane have an active Wnt signalling signature and are required for hair follicle formation, fibroblasts in the reticular dermal layer express high levels of genes associated with extracellular matrix (ECM) and immune signalling and mediate the initial phase of wound repair (7, 9, 11). During mouse skin development, dermal maturation is governed by a tight balance between fibroblast proliferation, quiescence and ECM deposition. Within the first week of postnatal life there is a coordinated switch in fibroblast behaviour from proliferative to quiescent, which is governed by ECM deposition/remodelling (12). While this quiescent state characterises postnatal skin, upon wounding, different fibroblast lineages are stimulated to proliferate and migrate into the wound site (13). Besides depositing/remodelling ECM in the wound bed, fibroblasts are able to acquire a dermal papilla or adipocyte fate in response to distinct signals and thereby promote hair follicle and DWAT regeneration, respectively (14–16). After tissue repair, the quiescent state of fibroblasts is restored.

UVB penetrates the epidermis and papillary dermis, while UVA affects the full thickness of the dermis, including the subcutaneous fat (5, 17). Photo-aged dermis is characterised by a loss of fibroblast density and changes in ECM organisation, including depletion of fibrillin-rich microfibers in the papillary dermis and accumulation of elastin-rich elastic fibres in the reticular dermis, which are mediated, at least in part by MMP activity (4, 5, 18, 19). In addition, UVR is a potent local and systemic immune modulator, able to modify the innate and adaptive immune response (3). Collectively, the activated signalling pathways and recruitment of distinct immune subsets lead to an immunosuppressive environment which supports inflammation resolution of a sunburn reaction and tissue repair but can also contribute to skin cancer.

While the consequences of UVR exposure for the epidermis, ECM and skin-resident immune network have been widely characterized (2, 3, 5), its short and long-term impact on different dermal fibroblast subpopulations (lineages) is unknown. In this study we have examined how dermal fibroblast lineages respond to acute and chronic UVB irradiation. Uncovering the UVR-induced early pathogenic processes leading to premature skin ageing and a cancer permissive environment will pave the way for new treatment strategies that target aberrant fibroblast behaviour (20).

## Results

### Acute UVR exposure results in a transient loss of fibroblasts in the papillary dermis

We began by determining the effect of acute UVB exposure on human dermal fibroblasts (21). For this study, 6 healthy volunteers (2 male, 4 female; mean age 44 ±12 years) were recruited. Their buttock skin was exposed to 3 times their individual minimal erythema dose (MED), sufficient to induce a moderate sunburn reaction characterised by histone H2AX phosphorylation (yH2AX), a central component of the DNA damage response and repair system, and cyclobutane pyrimidine dimers (CPD) accumulation in papillary fibroblasts (17). Skin biopsies were collected from irradiated skin at time-points up to 14 days post-UVR and subjected to double immunofluorescence labelling for CD39 (papillary fibroblast marker) and vimentin (VM, a pan-fibroblast marker) (11) (Figure 1A). Quantifying and plotting fibroblast density changes over time in skin sections of individual healthy volunteers revealed a pronounced loss of CD39/VM double positive cells in the upper dermis 1 and 4 days after UVR exposure (in 5 of 6 observed individuals) that returned to baseline levels after 10 or 14 days.

**Figure 1.**
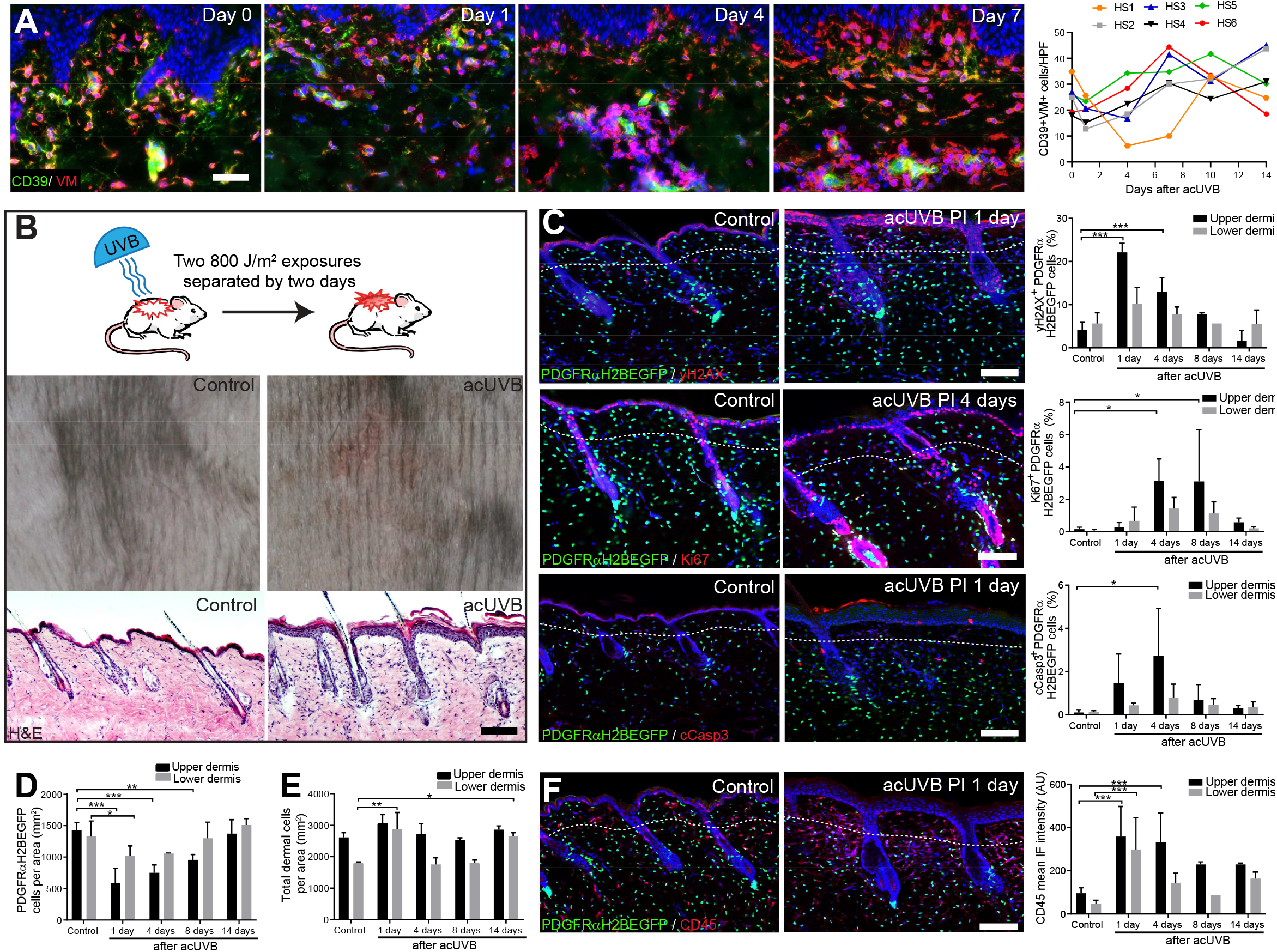
Acute UVB exposure depletes fibroblasts in the papillary dermis. (**A**) Immunostaining of human skin for CD39 (green) and VM (red) and quantification of double positive cells per field of view relative to control skin at indicated time points after acUVR exposure (n=6 biological replicates). (**B**) Experimental design of mouse acUVR model (top panel), representative images of skin erythema (middle panel) and H&E skin section (bottom panel), showing epidermal hyperplasia and increased dermal cell density 1 day after acUVR. (**C**) Representative PDGFRαH2BEGFP sections (green) stained for yH2AX (top panel), Ki67 (middle panel) and cCasp3 (bottom panel, red) of control and treated skin and quantification of double positive cells at indicated time points post-acUVR. Note that the epidermis and upper dermis show a pronounced DNA damage (yH2AX+) and proliferative response (Ki67+) with clusters of apoptotic cells (cCasp3+) 24 h post-acUVR (n=3-5 biological replicates). (**D, E**) Quantification of dermal fibroblast density (PDGFRαH2BEGFP+) (**D**) and total dermal density (DAPI+) (**E**) 24 h after acUVB in the upper and lower dermis (n= 3-7 biological replicates). (**F**) Immunostaining of PDGFRαH2BEGFP back skin (green) for all lymphocytes (CD45; red) and quantification of the CD45 mean fluorescence intensity at indicated time points post-UVR. Nuclei labelled with DAPI and dashed white line delineates upper and lower dermis. Scale bars, 50μm. Data are mean ±SD. *p<0.05,**p<0.01, ***p<0.001.

To uncover how different dermal fibroblast subpopulations are affected by UVB irradiation, we established an acute (ac)UVR mouse model consisting of two consecutive UVB exposures (800 J/m^2^) separated by 2 days, which induced moderate skin erythema – equivalent to a mild sunburn reaction in humans (Figure 1B) (6, 22, 23). Histology revealed epidermal hyperplasia, UVB-induced angiogenesis, increased dermal immune cell infiltration and temporary swelling of the papillary and reticular dermis at 1 day after UVB exposure (Figure 1B; Supplementary Figure 1, A and B). To study the effect of UVB on fibroblasts, we used the PDGFRαH2BEGFP transgenic mouse line, in which all dermal fibroblasts express nuclear GFP (24, 25) (Figure 1, C and D). Quantification of PDGFRαH2BEGFP positive cells showed a significant loss of upper dermal fibroblasts 1, 4 and 8 days after the second UVB exposure (Figure 1D), recapitulating the human in vivo findings (Figure 1A). The decrease in upper dermal fibroblasts correlated with an increase in DNA damage, measured by the phosphorylation of histone H2AX (yH2AX+), and apoptosis (cCasp3+) in dermal fibroblasts, particularly in the papillary dermis, 1 and 4 days post-UVB (Figure 1C). The number of immune cells (CD45+) increased markedly 1 day after irradiation and remained elevated for several days, accounting for the increase in total dermal cell density (DAPI+) during tissue repair (Figure 1, E and F; Supplementary Figure 1C). Fibroblast proliferation increased at day 4 and 8, particularly in papillary fibroblasts, returning to normal thereafter (Figure 1C, middle panel). No α-SMA+ dermal fibroblasts were present, indicating that UVB did not stimulate differentiation into myofibroblasts (Supplementary Figure 1D). We also noted that acUVB did not induce changes in the ECM that were detectable by CHP labelling, a molecular probe that recognises the triple helix structure of immature, damaged and remodelling interstitial collagen fibres (Supplementary Figure 1E) (26).

We conclude that in mouse and human skin, acute UVB exposure results in a transient loss of papillary fibroblasts, which is associated with dermal thickening, recruitment of immune cells and an increase in yH2AX and cCasp3 positive fibroblasts in the early acute UVB response and is followed by dermal fibroblast proliferation 4 days after UVB treatment.

### Chronic UVR induces long-term depletion of papillary fibroblasts and ECM changes

To test the impact of chronic UVR exposure on dermal fibroblasts we established a chronic (ch)UVR model consisting of 800 J/m^2^ UVB exposure twice a week for 8 weeks. This induced a prominent tanning response (melanin deposition) in UVR exposed back skin (Figure 2A) and a mild thickening of the epidermis, which correlated with an increase in Ki67 positive keratinocytes (Figure 2B). When the skin was examined 3 days after the final UVB treatment, there was no significant difference between control and UVB exposed dermis in terms of proliferation (Ki67+ fibroblasts) or apoptosis (cCasp3+ fibroblasts) and no α-SMA positive interfollicular fibroblasts were detected (Figure 2B; Supplementary Figure 2A). However, there was an increased abundance of CD45 positive cells in the upper and lower dermis and an increase in blood vessels (Figure 2B, lower panel; Supplementary Figure 2B). The ECM in chronically UVB exposed skin was highly remodelled. Herovici staining revealed accumulation of light blue stained immature collagen, particularly beneath the basement membrane following UVR; in contrast mature collagen in control skin stained pink/purple (Figure 2C). In addition, chUVR treated skin showed significantly increased CHP staining, indicating that collagen fibres were damaged or actively remodelled (Figure 2D) (26).

**Figure 2.**
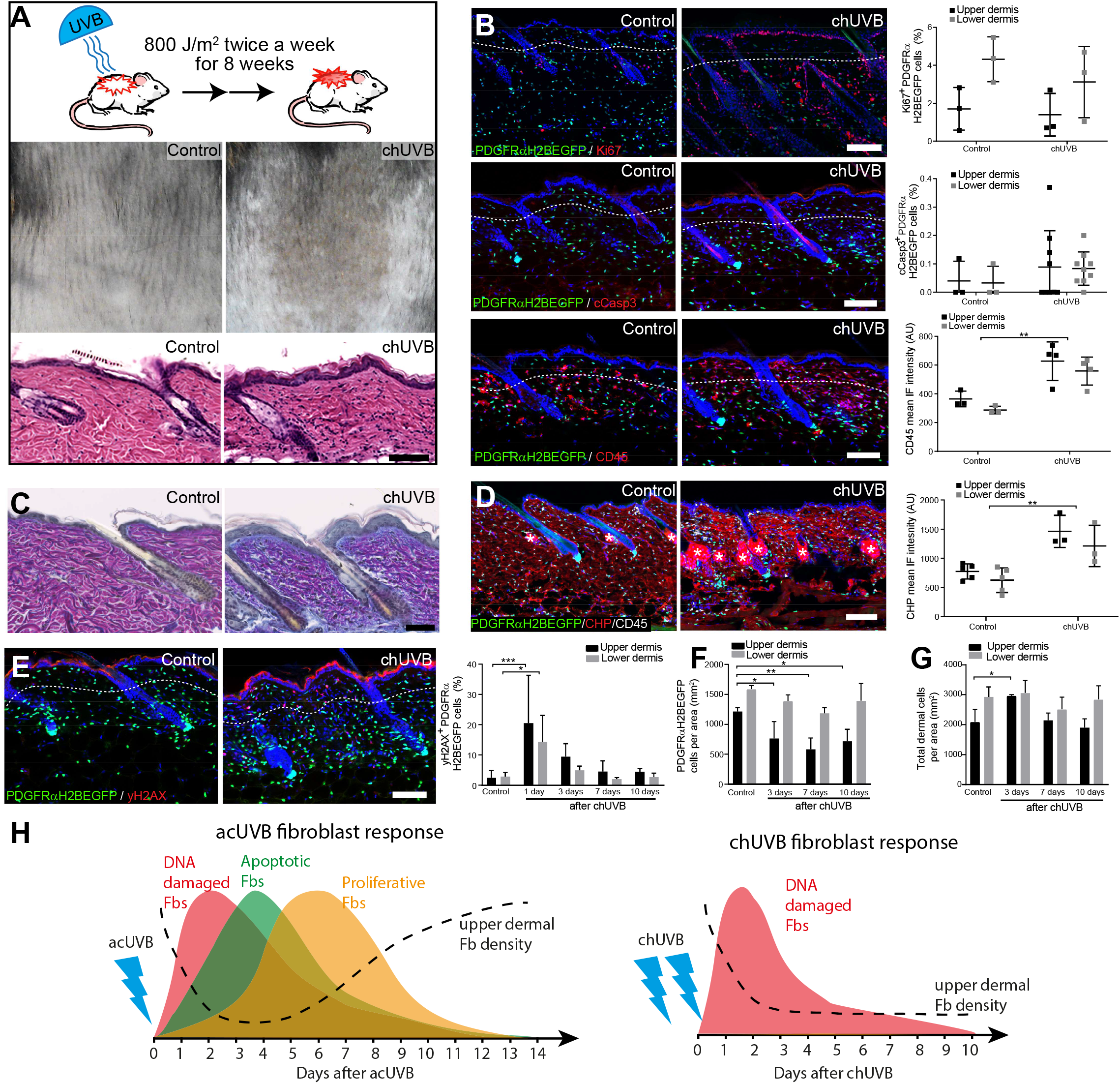
Chronic UVB irradiation leads to a permanent loss of papillary fibroblast in the upper dermis and changes in the ECM environment. (**A**) Experimental design (top panel), representative skin tanning (middle panel) and H&E section (bottom panel), showing epidermal hyperplasia, ECM changes and increased dermal cell density after chUVB. (**B**) Representative PDGFRαH2BEGFP sections (green) stained for Ki67 (top panel), cCasp3 (middle panel) and CD45 (bottom panel, red) of control and treated skin and quantification of either double positive cells (Ki67 and cCasp3) or mean fluorescence intensity (CD45) at 1 or 3 after last UVR exposure (n=3-5 biological replicates). While lymphocytes (CD45+ cells) are increased in the dermis, pronounced proliferation (Ki67+) and apoptosis (cCasp3+) are only observed in the epidermis after chUVB. (**C**) Herovici staining of control and chUVB exposed skin sections. Note that pink/purple staining indicates mature collagen, whereas light blue stained collagen in chUVB skin below the basement membrane is immature and actively remodelled. (**D**) Immunofluorescence staining of control and chUVB PDGFRαH2BEGFP skin (green) for CD45 (white) and collagen (red) using the CHP-biotin probe. Mean CHP fluorescence signal was quantified and increased CHP signal in chUVB skin indicates a more fibrillar, open and/or damaged collagen structure. White asterisks indicate unspecific CHP staining in sebaceous glands. (**E**) Immunostaining of control and chUVR exposed PDGFRαH2BEGFP back skin (green) for yH2AX (red) at 24 h after last UVB exposure and quantification of double positive cells at indicated time points (n=6 biological replicates). Note that the epidermis and dermis show a pronounced DNA damage (yH2AX+) at 24 h after UVR which is repaired over time. (**F, G**) Quantification of dermal fibroblast (PDGFRαH2BEGFP+) (**F**) and total dermal density (DAPI+) (**G**) after chUVB (n=3-4 biological replicates). (**H**) Comparison of acUVR and chUVR fibroblast tissue damage repair response. While acUVR induced a transient fibroblast depletion caused by DNA damage, fibroblast apoptosis and following proliferation, chUVR led to a persistent loss of fibroblasts in the papillary dermis. Nuclei were labelled with DAPI (blue) and dashed white line delineates upper and lower dermis. Scale bars, 50 μm. Data are mean ±SD. *p<0.05, **p<0.01, ***p<0.001.

chUVR induced significant DNA damage in the epidermis and dermal fibroblasts, which was observed 1 day after the final UVB exposure. The damage was progressively repaired post-UVR exposure, as measured by staining for yH2AX+ cells (Figure 2E). Quantification of fibroblasts (PDGFRαH2BEGFP+) in the upper and lower dermis showed that cell density was significantly decreased in the upper dermis after the final dose of UVR and was not restored to control levels even after 10 days (Figure 2F). In contrast, chUVR did not affect the density of fibroblasts in the lower dermis. Total dermal cell density (DAPI+) transiently increased at 3 days post-UVR, probably due to infiltrating immune cells (Figure 2, B and G).

We conclude that whereas acUVR leads to transient depletion of papillary fibroblasts, the effect is sustained after chronic treatment, correlating with more substantial ECM reorganisation. In contrast to acute UVR, there was minimal proliferation and apoptosis of dermal fibroblasts following chronic UVR (Figure 2H).

### Only fibroblasts in the papillary dermis contribute to repair of UVR damage

To understand how different fibroblast subpopulations contribute to regeneration of the papillary layer after acUVR exposure we performed lineage tracing (Figure 3A). Papillary fibroblasts can be specifically labelled with Lrig1CreER, while Dlk1CreER marks fibroblasts in the lower dermis when Cre-mediated recombination is induced at postnatal day 0 (P0) (7, 9). acUVR exposure of labelled transgenics confirmed the loss of papillary fibroblasts in the upper dermis, whereas Dlk1CreER labelled cells in the lower dermis were not affected (Figure 3B). At 4 days post-UVR papillary lineage dermal cells started to repopulate the upper dermis. However, the density of cells was significantly reduced compared to control dermis (Figure 3B). There were no detectable changes in the arrector pili muscle, dermal sheath and dermal papilla fibroblasts of hair follicles upon acUVR treatment (Figure 3B; Supplementary Figure 1D), suggesting that these fibroblasts did not contribute to repair of the damaged dermis (9, 27).

**Figure 3.**
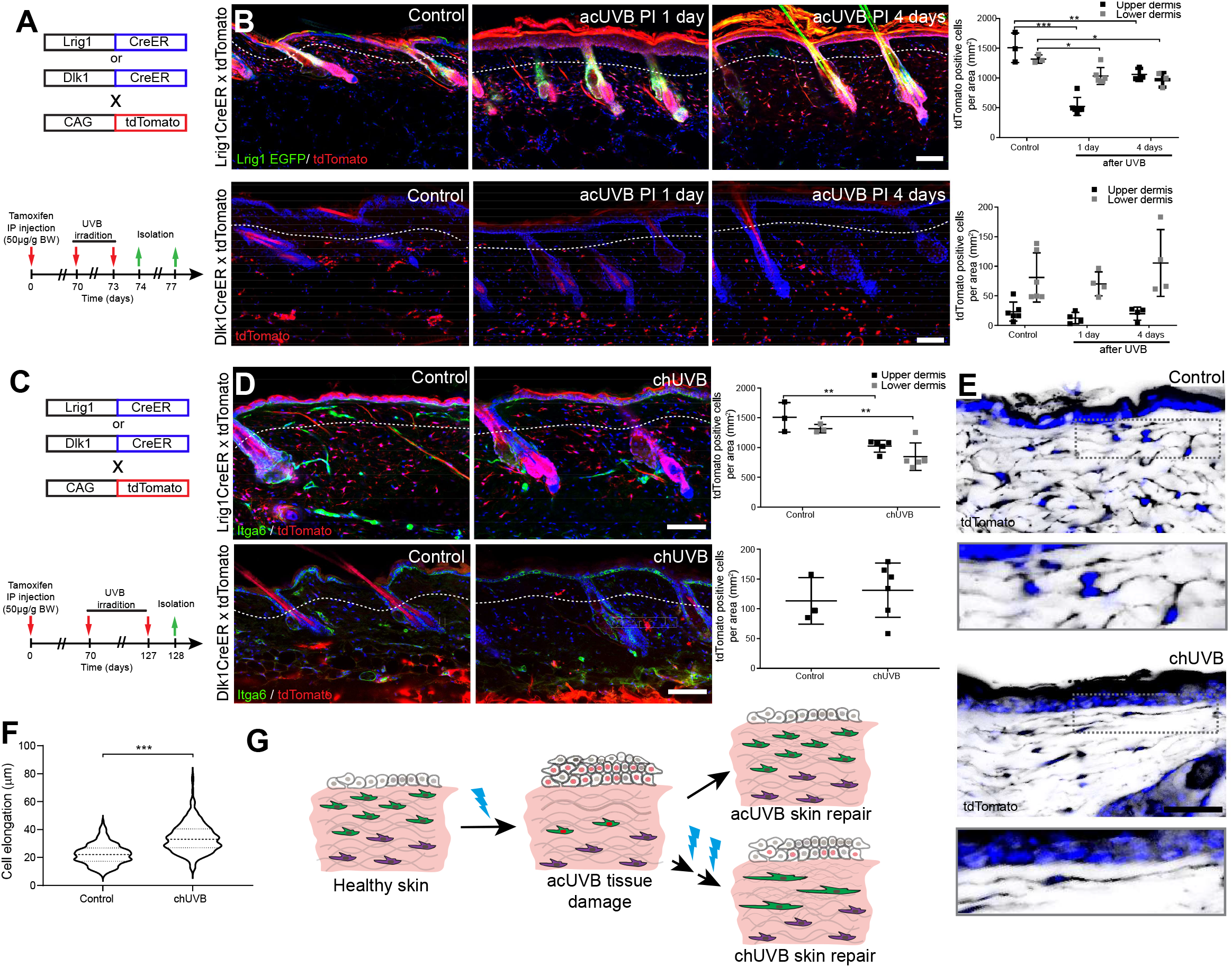
Only fibroblast lineages of the papillary dermis contribute to UVR induced tissue repair and fibroblasts in chronic UVR exposed skin are more elongated. (**A, B**) In vivo lineage tracing of distinct dermal fibroblast populations during tissue damage repair after acUVB. (**A**) Experimental design shows breeding strategy and skin isolation time points to follow fibroblast lineages during tissue repair. (**B**) Representative immunofluorescence image and quantification of Lrig1CreER x tdTomato (top panels) (n=3-6 biological replicates) and Dlk1CreER × tdTomato (lower panel) back skin of control and acUVB exposed skin after 1 and 4 days (n= 4-6 biological replicates). Quantification shows labelled cells in the upper and lower dermis at indicated time points. (**C, D**) In vivo lineage tracing of distinct dermal fibroblast populations during chUVR. (**C**) Experimental design shows breeding and lineage tracing strategy of chUVB exposed skin. (**D**) Immunofluorescence image and quantification of Lrig1CreER x tdTomato (top panels) (n= 3-5 biological replicates) and Dlk1CreER × tdTomato (lower panel) back skin of control and chUVB exposed skin 1 day after last UVR exposure (n= 3-6 biological replicates). Quantification shows labelled tdTomato+ cells in the upper and lower dermis. (**E, F**) Close up of Lrig1CreER x tdTomato lineage traced skin section showing cytoplasmatic tdTomato signal (black) (**E**) and quantification of papillary fibroblast elongation in control and chUVB exposed skin (**F**) (n= 300 cells from 4 biological replicates). Boxed areas in (**E**) indicate magnified fibroblasts shown below. Note that although fibroblasts density in chUVB skin is reduced (Figure 3D), fibroblast membrane protrusions are increased. (**G**) Summary of UVR-induced tissue damage and the skin regeneration after acute and prolonged (chronic) UVB exposure. In healthy skin papillary (green) and reticular (violet) fibroblast quiesce. After acUVR exposure papillary fibroblasts are depleted and epidermal and dermal cells start proliferating (red nucleus) during the tissue repair response. While fibroblast density and skin homeostasis are restored after acUVB tissue damage, repeated UVB exposure led to a permanent loss and elongation of papillary fibroblasts and changes in the ECM structure characteristic of aged skin. Nuclei were labelled with DAPI (blue) and dashed white line delineates upper and lower dermis. Scale bars, 50 μm. Data are mean ±SD. *p<0.05, **p<0.01, ***p<0.001.

Next, we investigated how chUVR exposure impacted the papillary (Lrig1CreER) and reticular (Dlk1CreER) fibroblast lineages (Figure 3C). As in the case of acUVR exposure Dlk1CreER labelled cells of the reticular dermis did not expand or contribute to tissue repair (Figure 3D). Lrig1CreER labelled cells were significantly reduced in the upper and lower dermis and showed a patchy distribution. Closer examination of papillary fibroblasts in chUVR skin revealed that, in contrast to acUVR exposure, their shapes were significantly elongated, suggesting that increased membrane protrusions may compensate for the fibroblast loss, as previously observed in aged skin(28) (Figure 3, E and F; Supplementary Figure 3).

We conclude that upon acUVR the upper dermis was replenished by papillary fibroblasts. In contrast, papillary fibroblasts in chUVR treated skin were not replenished and instead changed shape, increasing their cell membrane protrusions (Figure 3G). The lower dermal lineage was unaffected by acute or chronic UVR.

### Movement of papillary fibroblasts during the UVR tissue damage response

The number of fibroblasts in the papillary dermis is significantly lower in 1, 4 and 8 days post-acUVR skin than control (non-irradiated) skin (Figure 1D). To explore whether the papillary dermis is depleted via cell migration, we performed live imaging of anaesthetised PDGFRαH2BEGFP mice 1 day and 4 days after acUVR exposure (Figure 4A). In each case we recorded the movement of fibroblasts within defined fields up to 100 µm into the dermis, covering the papillary and upper reticular dermis, for 80 minutes. In agreement with previous measurements (12, 28), most fibroblasts in untreated adult skin maintained positional stability and showed minimal displacement over time (observed in 3 of 3 imaged biological replicates) (Supplementary Figure 4A; Supplementary Video 1).

**Figure 4.**
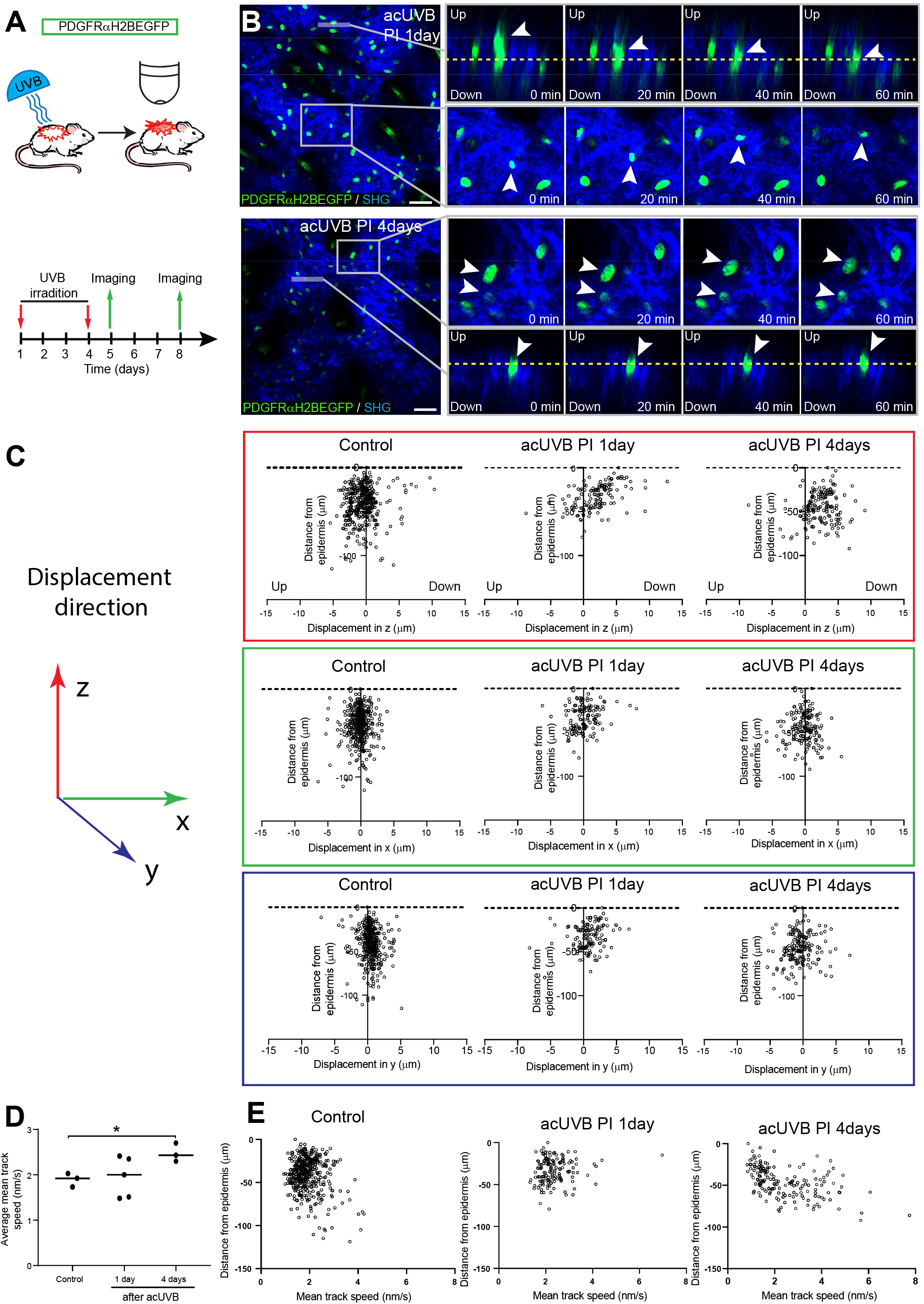
Fibroblasts in the papillary dermis become more motile during the UVR tissue repair response. (**A**) Experimental live imaging design of adult PDGFRαH2BEGFP back skin during acUVB induced tissue damage repair. (**B**) Representative time-lapse images of adult PDGFRαH2BEGFP (green) dermis 1 days (upper panel, relates to Supplementary Video 2) and 4 days (lower panel, relates to Supplementary Video 3) post-acUVB with collagen shown as second harmonic generation (SHG) in blue at indicated imaging time points. Line indicates orthogonal close up to follow vertical cell displacement and box shows fibroblast movement in the horizontal plane. Arrow heads in close ups indicate cells migrating and dashed line is for orientation. (**C**) Scatter plots of the displacement along the indicated axis (z-, red; x-, green; y-axis, blue) of individual control and acUVB treated cells in their relative z-location (distance from epidermis). (**D**) Average mean cell displacement speed of imaged control and acUVB exposed back after 1 and 4 days (n=3-5 biological replicates). (**E**) Scatter plots of mean velocity of individual cells in their relative z location from representative control and acUVB treated animals after 1 and 4 days post-UVB. Scale bars, 50 μm. *p<0.05.

One day after UVR single papillary fibroblasts could be observed for example moving into the deeper dermis or within the horizontal plane (Figure 4B, top panel; Supplementary Video 2). Quantification of cell displacement in the horizontal and vertical directions indicated that fibroblasts start to be more motile along the (z) axis 1 day post-irradiation compared to control skin (Figure 4C; Supplementary Figure 4B). However, most fibroblasts displayed minimal displacement and the direction of the observed movement appeared heterogenous (Figure 4C; Supplementary Figure 4B). At 4 days after UVR exposure more fibroblasts showed increased random cell displacement within the horizontal and vertical dermal plane across the imaged dermis (Figure 4, B and C; Supplementary Figure 4B; Supplementary Video 3). In line the average mean displacement speed of fibroblasts was more heterogenous at 1 day post-UVR compared to control skin and increased significantly at 4 days post-UVR (Figure 4D). Plotting the individual cell mean speed across the upper dermis revealed that fibroblasts with increased motility were present throughout the upper dermis at 4 days after UVR (Figure 4E).

We conclude that fibroblast depletion in the papillary layer in the early UVR response is associated with minimal migration, whereas at 4 days post-acUVR fibroblasts become more motile, which correlates with ECM remodelling and fibroblast redistribution (Figure 3, B and G). The lack of directional migration indicates that fibroblast replenishment of the papillary dermis after UVR damage is a stochastic process similar to the fibroblast lineage redistribution observed during dermal maturation and ageing (12).

### Activation of Wnt signalling does not enhance dermal recovery after UVR exposure

Wnt/β-catenin signalling plays a major role in skin development, wound healing, regeneration and cancer (10, 29). To explore how Wnt/β-catenin signalling is regulated by UVR we subjected TOPEGFP reporter mice to acUVR. In these mice H2BeGFP is expressed under the control of multiple Lef1/TCF binding sites, allowing nuclear GFP expression to be used as a readout of Wnt/β-catenin signalling activity (30). Wnt/β- catenin activity was highly induced in epidermal and dermal cells at 1 and 4 days post-acUVR (Figure 5A), which coincided with fibroblast DNA damage repair and proliferation (Figure 1C). During tissue repair and fibrosis Wnt signalling has been shown to cooperate with YAP/TAZ signalling at multiple levels (31, 32). Consistent with this, dermal fibroblasts of the papillary dermis and IFE keratinocytes displayed increased nuclear YAP localisation in acute and chronic UVR exposed skin (Figure 5B; Supplementary Figure 5).

**Figure 5.**
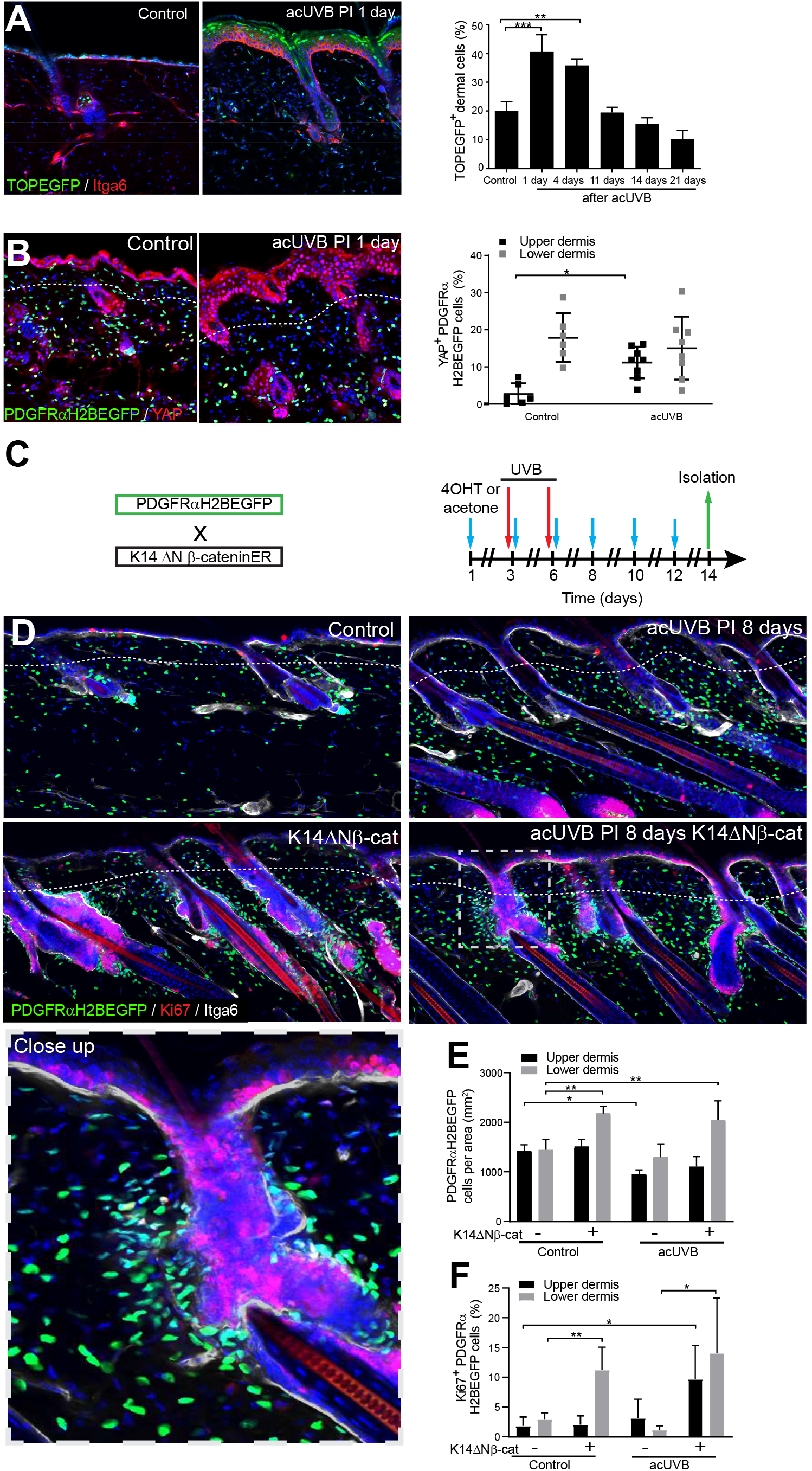
Induction of fibroblast proliferation is not sufficient to restore dermal homeostasis after UVR exposure. (**A**) Representative Wnt signalling reporter (TOPEGFP) section stained for Itga6 (red) of control and treated skin, in which H2BEGFP (green) is expressed under the control of multiple Lef1/TCF binding sites reporting active Wnt/β-catenin signalling(30) and quantification of TOPEGFP positive dermal cells in control and UVR treated skin (n=3-5 biological replicates). Note that Wnt/β-catenin signalling is increased in the epidermis as well. (**B**) Representative PDGFRαH2BEGFP back skin sections (green) for YAP (red) 1 day after acUVB exposure and quantification of PDGFRαH2BEGFP positive cells with nuclear YAP in the upper and lower dermis (n=6-8 biological replicates). Nuclear YAP is increased in the papillary dermis and IFE after acUVB exposure. (**C**) Experimental strategy for increasing fibroblast proliferation during acUVB damage tissue repair by stabilizing epidermal β-catenin (K14ΔNβ-cat transgenic). (**D**) Representative PDGFRαH2BEGFP back skin sections (green) of indicated transgenic stained for Ki67 (red) after isolation at 8 days post-UVR. Dashed box indicates close up area shown in the lower panel. (**E, F**) Quantification of dermal fibroblast density (PDGFRαH2BEGFP+) (**E**) and proliferation (Ki67+ PDGFRαH2BEGFP cells) (**F**) at indicated treatment conditions (n=3-8 biological replicates). Nuclei were labelled with DAPI (blue) and dashed white line delineates upper and lower dermis. Scale bars, 50 μm. Data are mean ±SD. *p<0.05,**p<0.01,***p<0.001.

To test whether induction of fibroblast proliferation by epidermal Wnt signalling modified the response to UVR, we crossed PDGFRαH2BEGFP mice with K14ΔNβ-cateninER mice, which express stabilized β-catenin under the control of the K14-promoter in response to Tamoxifen application. This has previously been shown to promote fibroblast proliferation and ECM remodelling in the absence of dermal inflammation (Figure 5C) (33). Analysing fibroblast distribution 8 days post-UVR and epidermal β-catenin stabilisation in PDGFRαH2BEGFP x K14ΔNβ-cateninER transgenics revealed that fibroblasts predominantly proliferated and expanded around existing and ectopic hair follicles in the lower dermis (Figure 5, D, E and F). However, this increased abundance of fibroblasts failed to efficiently repopulate the interfollicular dermis beneath the basement membrane and failed to restore fibroblast homeostasis and organisation in the papillary dermis.

In conclusion, although Wnt signalling is activated by UVR, increasing fibroblast proliferation by genetically stabilizing epidermal β-catenin was not sufficient to improve fibroblast regeneration of the papillary dermis.

### Dermal fibroblast survival is supported by cutaneous T cells controlling the inflammatory response to UVR exposure

The inflammatory response to UVR is well documented. Neutrophils are the first immune cell type to infiltrate into the dermal region after UVB exposure (34, 35) and this is followed by an influx of different T cell populations (3). In our acUVR model we observed an increase in CD45 + cells that persisted even after 14 days (Figure 1F; Supplementary Figure 1C). The number of neutrophils increased in skin 1 day post-UVR (Supplementary Figure 6, A and B). This was followed by an increase in the abundance, proliferation and activation of different T cell populations, specifically CD8+ cytotoxic T cells and FoxP3+ regulatory T cells (Tregs). Immunofluorescence analysis of CD3, CD8 and FoxP3 labelling revealed that while CD3+ T cells were mainly depleted in the epidermis, cytotoxic T cells and Tregs expanded throughout the dermis at 3 days after acUVR exposure (Supplementary Figure 6C). This is in line with the observed immune cell behaviour in human skin upon UVB exposure where neutrophil infiltration is followed by an increase in accumulation, activation and proliferation of different T cell populations (21, 36, 37).

**Figure 6.**
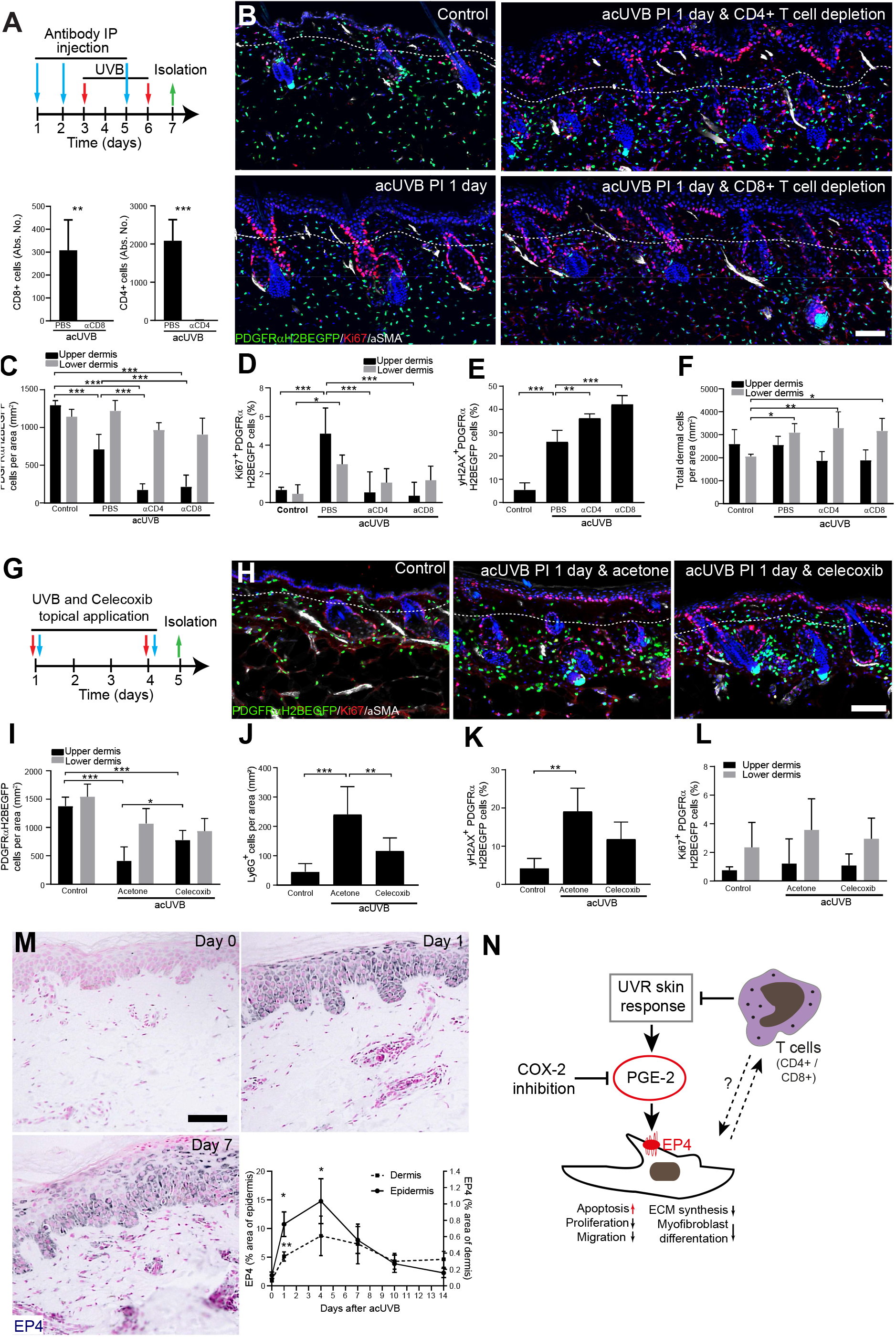
Papillary fibroblast survival is dependent on cutaneous T cells controlling the inflammatory response to UVR exposure. (**A**-**F**) CD4 and CD8 positive cell depletion increased fibroblast loss in the upper dermis after acUVB. (**A**) Experimental strategy for antibody based immune cell depletion during acUVB (blue arrow, antibody injection; red arrow, UVB; green arrow, skin isolation) (top panel). Antibody depletion was assessed by FACS analysis of cutaneous CD4 and CD8 positive cells and absolute number quantifications are for 6 cm^2^ (bottom panels) (n=4 biological replicates). (**B**) Representative immunostaining of PDGFRαH2BEGFP back skin (green) for Ki67 (red) and α-SMA (white). (**C-F**) Quantification of dermal fibroblast density (PDGFRαH2BEGFP+) (**C**), proliferation (Ki67+ PDGFRαH2BEGFP cells) (**D**), DNA damage (yH2AX+ PDGFRαH2BEGFP cells) (**E**) and total dermal density (DAPI+) (**F**) 24 h after acUVB (n=4 biological replicates). (**G-L**) Inhibition of UVR induced inflammation decreased fibroblast loss. (**G**) Experimental design for topical treatment with Celecoxib (COX-2 inhibition) immediately after acUVB exposure. (**H**) Representative immunostaining of PDGFRαH2BEGFP back skin (green) for Ki67 (red) and α-SMA (white) at indicated treatment condition. (**I-L**) Quantification of dermal fibroblast density (PDGFRαH2BEGFP+) (**I**), neutrophil infiltration (Ly6G+) (**J**), DNA damage (yH2AX+ PDGFRαH2BEGFP cells) (**K**) and fibroblast proliferation (Ki67+ PDGFRαH2BEGFP) (**L**) at indicated experimental condition (n=4-8 biological replicates). (**M**) Immunostaining of human skin for EP4 receptor and quantification of EP4 in the epidermal and dermal area per field of view at indicated time points after acUVB exposure (n= 13 biological replicates). (**N**) Model of PGE-2-EP4 signalling in dermal fibroblasts after UVR exposure influencing their tissue damage response and survival in concert with other T cells. Data are mean ±SD except in (M) that is ±SEM. *p<0.05, **p<0.01, ***p<0.001. Nuclei were labelled with DAPI (blue) and dashed white line delineates upper and lower dermis. Scale bars, 50 μm. IP, intraperitoneal injection.

To explore the functional consequences of the increase in different T cell populations after UVR, we depleted CD4+ and CD8+ cells with specific blocking antibodies before and during acUVB exposure (Figure 6A) (38–40). Back skin from control and UVR treated mice was isolated 24 h after the last treatment and flow cytometric analysis of skin and lymph nodes confirmed successful immune cell depletion (Figure 6A; Supplementary Figure 6, D and E). Depletion of either CD8+ or CD4+ cells significantly increased the loss of upper (papillary) fibroblasts and was associated with a significant increase in DNA damage (yH2AX+ fibroblasts) and a reduction in fibroblast proliferation (Ki67+) (Figure 6, B, C,D and E). Elevated proliferation in the basal layer of the epidermis upon acUVB exposure was not affected by the blocking antibody treatments (Figure 6B; Supplementary Figure 6F). Although total dermal cells (DAPI+) in the lower dermis were significantly increased, PDGFRαH2BEGFP positive cells were unchanged in the reticular dermal layer (Figure 6, C and F).

Previous reports have suggested that CD4+ T cell depletion significantly increases and prolongs the acute UVB-induced cutaneous inflammatory response (40). To test if fibroblast depletion was mainly caused by an UVR-induced inflammatory response, we treated back skin topically with the COX-2 inhibitor celecoxib immediately after UVR exposure (Figure 6G). Topical treatment with celecoxib following UVB irradiation has been shown to inhibit several parameters of UVR induced acute inflammation, including vascular permeability, infiltration of neutrophils, prostaglandin E_2_ (PGE-2) production as well as acute oxidative damage (41, 42). Notably, COX-2 inhibition significantly inhibited fibroblast depletion in the upper dermis and neutrophil infiltration (Ly6G+) after UVR treatment (Figure 6, H, I and J). Furthermore, DNA damage in dermal fibroblasts (yH2AX+) was reduced (Figure 6K). In contrast celecoxib treatment did not affect fibroblast proliferation (Ki67+) and total dermal cell density (DAPI+) after acute UVR exposure (Figure 6L; Supplementary Figure 6G).

It has been reported previously that following acUVB exposure of human skin pro-inflammatory prostaglandins including PGE-2 peak at 24 h before normalising after 4 days (21, 37). Besides inhibiting fibroblast proliferation, collagen synthesis, migration and differentiation into myofibroblasts, PGE-2 increases fibroblast apoptosis through E prostanoid (EP)2 and EP4 receptor signalling, resulting in activation of phosphatase and tensin homologue on chromosome 10 (PTEN) and downstream inhibition of protein kinase B /AKT, an important pro-survival signal (43). EP4 is the major EP receptor in skin (44) and is expressed by dermal fibroblasts (45, 46). Immunostaining skin sections for EP4 from UVB-treated humans revealed that EP4 expression was strongly increased in the epidermis and dermis at 1 and 4 days after acUVB exposure before returning to lower levels after 7 days in both skin compartments (Figure 6M). These findings suggest that the UVB-induced increase in PGE-2 level and EP4 expression influences the dermal fibroblast UVB damage response and survival (Figure 6N).

In summary we have shown that the loss of papillary fibroblasts in mice is primarily induced by an UVR induced inflammatory response that can be supressed by COX-2 inhibition. Infiltrating/activated T cells support fibroblast survival and regeneration following UVR induced environmental stress (Figure 6N). Studies of UVR treated human skin implicate a role for increased PGE-2 production.

## Discussion

In this study we have elucidated the short- and long-term impacts of UVR exposure on different fibroblast lineages in the skin. We reveal that physiological doses of UVR are sufficient to severely deplete papillary fibroblasts in human and mouse skin, and that fibroblast survival is influenced by cutaneous T cells and PGE-2/EP4 receptor signalling. Our immunofluorescence, lineage tracing and in vivo live imaging showed that the loss of papillary fibroblasts is primarily due to apoptosis rather than movement of papillary fibroblasts into the deeper dermis. After acute UVR fibroblasts start proliferating, increase motility and restore tissue density. In contrast prolonged exposure to UVR prevented repopulation of fibroblasts in the upper dermis even after 10 days post-UVR (Figure 2H). Loss of the papillary lineage is associated with premature skin ageing, reduced regeneration and a profibrotic environment (7, 9, 33, 47).

We and others have recently shown how different fibroblast lineages contribute to dermal architecture and have explored their tissue-scale behaviour in development and skin regeneration (12, 28, 48). Deregulation of these complex processes is associated with several skin pathologies, including fibrosis, chronic wounds and cancer. Comparison of the fibroblast lineage response during repair of full thickness wounds and UVR-induced tissue damage reveals several differences. While both forms of tissue damage induce a pronounced inflammatory response and activation of Wnt/β-catenin and YAP/TAZ signalling in the epidermis and dermis, UVR-induced tissue damage is repaired with minimal fibroblast proliferation and myofibroblast differentiation. We recently showed that ECM is a potent regulator of fibroblast behaviour and inhibits proliferation during skin development and regeneration (12). The inhibitory signal of the ECM can be partly overcome by overexpression of epidermal β-catenin, which induces the expression of several fibroblast growth factors (33). However, the induction of proliferation was not sufficient to restore fibroblast organisation in UVR damaged skin.

Our lineage tracing and in vivo live imaging experiments revealed that only fibroblasts in the upper dermis contribute to tissue repair (Figure 3G). While in the early UVR response (1day post-UVR) fibroblast migration is limited to single cells, papillary fibroblasts become more motile at 4 days post-UVR, which could be due to ECM remodelling. In support of this concept, during skin homeostasis most dermal fibroblasts are stationary, yet active random fibroblast migration has been observed close to growing hair follicles where the surrounding ECM is extensively remodelled (28). In contrast, upon full thickness wounding we and others have shown that fibroblasts start migrating towards the wound where they randomly distribute and expand during the early wound repair phase (12, 49). During wound healing chemo-attractants such as platelet-derived growth factor (PDGF) are key regulators of fibroblast chemotaxis (50); however, the intrinsic and extrinsic signals controlling fibroblast migration after UVR exposure remain unclear.

Recent laser or genetic ablation experiments have revealed that loss of dermal fibroblasts is repaired through a mixture of proliferation/migration and reorganisation of the plasma membrane network(28). Our data indicate that a similar mechanism may apply during repair of UVR induced tissue damage. In contrast to acUVR tissue damage, in chUVR skin the decreased fibroblast density persisted in the papillary dermis and surviving fibroblasts were significantly elongated, suggesting that in photoaged skin, fibroblast loss is also compensated by an increased membrane network of surrounding fibroblasts. This is in line with the observation of Marsh *et al*. that the progressive loss of fibroblasts during skin ageing is balanced by increasing membrane protrusions rather than fibroblast proliferation or migration (28).

Resident immune cells are not only essential for skin barrier function, pathogen defence and wound healing but also provide essential signals for hair follicle growth and skin regeneration (14, 38, 51). Here we identify an additional function, that of promoting the survival of dermal fibroblasts during environmental stress. In line with previously published reports, we found that upon UVR exposure neutrophils were first recruited to the UV exposed site (21, 42, 52). This was followed by infiltration of different types of T cells; in particular Tregs became highly activated and proliferative. While in homeostatic conditions Tregs are predominantly located around hair follicles (38), their spatial expansion throughout the interfollicular dermis was evident upon UVR exposure, and this could potentially promote an immunosuppressive environment. Whether there is also direct cross talk between T cells and dermal fibroblasts during the UVR tissue damage response is currently unclear.

A recent study in human skin has identified CD4+ GATA3+ and CD8+ GATA3+ T cells as the predominant T cell populations in UVR induced inflammation and these are therefore likely to contribute to tissue resolution *via* dermal communication (21). Neutrophil derived reactive oxidants are potent mediators of UVB-induced tissue damage and tumorigenesis because of their cytotoxicity and immunosuppression (34, 35). Production of multiple pro-inflammatory prostaglandins, including PGE-2, is promoted following UVR as a result of arachidonic acid release by phospholipases and by induction of COX-2 expression in various skin cells (37, 53, 54). PGE-2 governs diverse biological functions that are mediated by signalling through four distinct E-type prostanoid (EP) receptors, EP1-4 (55). The major EP receptor in skin is EP4, which is a Gs coupled receptor regulating cAMP/PKA, MEK/ERK1/2, NF-кB and PI3K/ERK/Akt signalling; these pathways are important for cell survival, proliferation, migration, differentiation, angiogenesis and inflammation. In fibroblasts PGE-2 binding to EP4 has been shown to increase PTEN activity and Fas expression and decrease survivin expression, thereby promoting apoptosis (43). In line with these observations, the increased PGE-2 levels and EP4 expression in the dermis in the early UVR response coincide with the observed fibroblast depletion at 1 and 4 days after UVB exposure (Figure 6N).

In support of the concept that prostaglandin signalling influences fibroblast survival, specific inhibition of COX-2 immediately after UVB exposure significantly reduced fibroblast depletion and neutrophil recruitment. This is consistent with the observation that the COX enzyme inhibitor aspirin (acetylsalicylic acid) efficiently protects keratinocytes and melanocytes from acute UVB-induced DNA damage by decreasing cutaneous inflammation and PGE-2 levels in skin (56). Based on our findings, aspirin may also enhance dermal fibroblast survival and regeneration upon UVR induced environmental stress. Whether the anti-inflammatory activities of aspirin could help prevent fibroblast associated changes in photo-aged skin warrants investigation in the future.

In summary we have shown that papillary fibroblasts repair the dermis following UVR and that their survival is influenced by tissue resident T cells. We have previously reported that papillary fibroblasts differ from reticular fibroblasts in expressing genes related to Wnt, ECM and inflammation (11). This may explain their unique responses to UVR.

## Methods

### Human volunteer and UVR time course analysis

Ethical approval was granted by the Greater Manchester North NHS research ethics committee (ref: 11/NW/0567) for the studies presented in Figure 1 and Figure 6. Details of the time course analysis of UVR challenged human skin have been reported previously (21). Briefly, this study was conducted at the Photobiology Unit, Salford Royal NHS Foundation Trust, Greater Manchester, UK and involved healthy volunteers aged between 18 and 60 years. All were white Caucasian, of skin phototypes I-III according to the Fitzpatrick skin phototyping scale. In each case photo-protected upper buttock skin received 3 times the minimal erythema dose (MED) using a UVB lamp (Waldmann 236B, peak 313 nm, 280-400 nm) to separate sites on five different days. This allowed collection of skin samples at 1, 4, 7, 10 and 14 days post-UVR, in addition to unirradiated skin. Erythema measurements and 5 mm skin punch biopsies were taken at the end of the time course experiment, as described (21). Skin biopsies were bisected, with half snap frozen in optimal cutting medium and half formalin fixed and paraffin embedded. All volunteers provided written informed consent in accordance with the Declaration of Helsinki principles.

### Transgenic mice

All animal experiments were subject to local ethical approval and performed under the terms of a UK government Home Office license (PPL 70/8474 or PP0313918). All mice were outbred on a C57BL/6 background and male and female mice were used in experiments that included PDGFRαH2BEGFP(24), Lrig1CreERt2-IRES-GFP (Lrig1CreER) (57), Dlk1CreERt (Dlk1CreER) (7), K14ΔNβ-cateninER (K14ΔNβ-cat)(58), ROSAfl-stopfl-tdTomato (Jackson Laboratories, 007905) and TopH2BeGFP (TOPEGFP) (30) mice. Animals were sacrificed by CO_2_ asphyxiation or cervical dislocation. All efforts were made to minimise suffering for mice. For lineage tracing, transgenic reporter mice were crossed with indicated CreER line which was induced by injection with 10 µl tamoxifen (50 µg/g body weight; Sigma-Aldrich) intraperitoneally in newborn mice (P0), when Dlk1 and Lrig1 are highly expressed in dermal fibroblasts (7, 9). Tamoxifen for injection was dissolved in corn oil (5 mg/ml) by intermittent sonication at 37°C for 30 min. For epidermal β-catenin stabilisation acUVR experiments, central back skin of K14ΔNβ-cat x PDGFRαH2BEGFP transgenics was clipped and treated topically with 100 µl 4-Hydroxytamoxifen (4OHT) (2 mg/ml dissolved in acetone; Sigma-Aldrich) every second or third day for a total of 6 applications before and after sham or acUVB exposure (see experimental design in Figure 5C). Tissue was collected at the indicated time points, briefly fixed with 4% paraformaldehyde/PBS (10 min at room temperature) and embedded into optimal cutting temperature (OCT) compound or fixed 4% paraformaldehyde/PBS overnight at 4°C for paraffin embedding as previously described (59).

### UVR acute and chronic mouse models including COX-2 inhibition and immune cell depletion

For the in vivo UVR treatments, a UVR system (Tyler Research UV-2) was used which has a cascade-phosphor UV generator lamps with a sharp 310 nm peak output (65% of UVR falls within 20 nm half bandwidth). Thus the generated UVR is highly enriched for UVB which penetrates the epidermis and upper dermis (5). Comparison of different mouse strains has revealed that the used C57BL/6 background most closely mimics the human skin UVB response (23, 60). For the UVR exposure, mice were restrained in a custom-made mouse restrainer only exposing a defined (2 cm x 3 cm) central back skin area to the UVR. The UVB dose used for the acute (two consecutive exposures separated by two days and isolation at indicated time points after second treatment) and chronic (twice a week for 8 weeks and isolation at indicated time point after last exposure) models (800 J/m^2^) has been shown to closely correspond to the clinically relevant UVB dose in C57BL/6 mice inducing a detectable skin reaction (erythema/oedema) (23).

For immune cell depletion in vivo anti-CD4 (clone GK1.5, 400 mg per injection in 100 µl PBS) and anti-CD8 (clone 2.43, 400 mg per injection in 100 µl PBS) both purchased from BioXcell (West Lebanon, NH, USA) were administered intraperitoneally 3 times before and during acUVR exposure (see experimental design Figure 6A). Back skin and lymph nodes were collected 24 h after the last treatment and analysed by flow cytometry and immunofluorescence.

For the COX-2 inhibition during acUVR, mice were divided into control and UVR treated groups which were either treated topically with vehicle (200 µl acetone) or 500 µg of celecoxib (Selleckchem, S1261) dissolved in acetone (200 µl) immediately after sham or UVR exposure (800 J/m^2^) (see experimental design Figure 6G). Mice were killed 24 h after the second UVB treatment and back skin was analysed as described above.

In all UVR experiments 10-20 weeks old male and female mice were randomised in the different experimental groups and the hair of the central back skin was clipped 24 h prior to sham or UVR exposure. Back skin with hair follicles not in telogen (hair growth resting phase) at the beginning of the experiment was excluded. During the UVR procedures mice were housed in small groups (≤3) to minimize the risk of fighting, and skin with signs of scratching was not considered in the analysis.

### Tissue digestion and flow cytometry analysis

Preparation of single cell suspensions for flow cytometry was performed as previously described (38). Briefly, isolation of cells from skin draining lymph nodes (axillary, brachial and inguinal lymph nodes) for flow cytometry was performed by mashing tissue over 70 µm sterile filters. For isolation of skin cells, mouse dorsal skin was minced finely, re-suspended in 3 ml of digestion mix (composed of 2 mg/ml collagenase XI, 0.5 mg/ml hyaluronidase and 0.1 mg/ml DNase in 10% Fetal Bovine Serum, 1% Pen/Strep, 1 mM Na-pyruvate, 1% HEPES, 1% non-essential amino acid, 0.5% 2-mercaptoethanol in RPMI-1640 (+L-glut) medium) and digested for 45 min at 37°C 255 rpm. The digestion mix was then resuspended in 20 ml of RPMI/HEPES/P-S/FCS media and passed through a 100 µm and 40 µm cell strainer before centrifugation at 1800 rpm for 4 min at 4°C. Cell pellets were resuspended in 1 ml of FACS buffer (2% Fetal calf serum, 1 mM EDTA in PBS) for cell counting with an automated cell counter (NucleoCounter NC-200, Chemometec) to calculate absolute cell numbers. Following isolation from the tissue, cells were stained with surface antibodies and a live dead marker (Ghost DyeTM Violet 510) (see Supplementary Table1) for 20 min on ice. All samples were run on Fortessa 2 (BD Bioscience) in KCL BRC Flow Cytometry Core, which was standardized using SPHERO Rainbow calibration particle, 8 peaks (BD Bioscience, 559123). For compensation, UltraComp eBeads™ (Thermofisher, 01-2222-42) were stained each surface and intracellular antibody following the same procedure as cell staining. ArC™ Amine Reactive Compensation Bead Kit (Thermofisher, A10346) were used for GhostDye™ Live/Dead stain. All gating and data analysis were performed using FlowJo v10, while statistics were calculated using Graphpad Prism 9.

### Histochemical and Immunostaining

H&E and Herovici staining of 8 μm thick paraffin mouse skin sections were processed as previously described (59) and sections were mounted in DPX mounting medium (Sigma-Aldrich).

For immunostaining, mouse tissue samples were embedded in optimal cutting temperature compound (OCT, Life Technologies) prior to sectioning. For thin section stains, cryosections of 14 µm thickness were fixed with 4% paraformaldehyde/PBS (10 min at room temperature), permeabilised with 0.1% Triton X-100/PBS (10 min at room temperature), blocked with 5% BSA/PBS (1 h at room temperature) and stained with the following primary antibodies to: vimentin (1:500; Cell Signaling, #5741), Ki67 (1:500; Abcam, ab16667 and Invitrogen, clone SolA15), α-SMA (1:500; Abcam, ab5694), Krt14 (1:1000; BioLegend, 906001), yH2AX (1:500; Abcam, ab81299 and), YAP (1:500; Cell Signaling, #14074), cCasp3 (1:500; Cell Signaling, #9661), CD45 (1:200; eBioscience, clone 30-F11), CD31 (1:200; eBioscience, clone 390), CD49f (1:500; BioLegend, clone GoH3), CD3 (1:200; BioLegend, clone 17A2), CD8 (1:200; BioLegend, clone 53-6.7), FoxP3 (1:200; eBioscience, clone FJK-16s) and Ly6G (1:200; eBioscience, clone 1A8). Samples were stained overnight at 4°C, washed in PBS, labelled with secondary antibodies (all 1:500; AlexaFluor488, A-21208; AlexaFluor555, A-31572; AlexaFluor555, A-21434; AlexaFluor647, A-21247; Thermo Fisher) for 1 h at room temperature and stained for 10 min with 4,6-diamidino-2-phenylindole (DAPI; 1 mg/ml stock solution diluted 1:50,000 in PBS; Thermo Fisher) at room temperature with at least four PBS washes in-between. For horizontal wholemount, 60 µm sections were immunostained as described previously(9). While thin sections were mounted with ProLong Gold Antifade Mountant (Thermo Fisher), horizontal wholemounts were mounted with glycerol.

For human tissue fibroblast stainings, cryosections of 7 µm thickness were fixed with 4% paraformaldehyde/PBS (20 min at room temperature), blocked with 2.5% normal horse serum/TBS (20 min at room temperature) and then incubated with the following primary antibodies: CD39 (eBioscience, clone eBioA1 (A1)), in blocking solution; 1:200) and vimentin (Cell Signaling, #5741; 1:500) overnight. After three TBS washes sections were incubated for 30 min with anti-mouse A488 and anti-rabbit A594 secondary antibodies (VectorFluor Dylight Duet kit; DK-8828). Thereafter sections were incubated with DAPI, washed and mounted. For the EP4 immunostaining in human skin, paraffin sections of 5μm thickness were rehydrated, permeabilised with 0.5% Triton X-100/TBS (10 min) and endogenous hydrogen peroxide activity was blocked using 0.3% hydrogen peroxide/PBS (10 min). After blocking with 2.5% normal horse serum/TBS tissue sections were incubated with rabbit polyclonal EP4 (1:50; Cayman, Ann Arbor, Michigan, USA) for 1 h at room temperature. Primary antibody binding was visualised using a Vector ImmPress kit (Vector Labs, Peterborough, UK) according to the manufacturer’s instructions, and sections counterstained with nuclear fast red (Vector labs). Images were acquired using the 3D Histech Pannoramic 250 Flash II slide scanner with a (20x/0.80 Plan Apo) objective. EP4 expression was analysed in 3 skin sections (3 fields of view per section) per time point using Image J software (NIH, UK); thresholding was used to mask positively stained areas and the percentage area of epidermis or dermis was calculated.

For collagen hybridising peptide (CHP) staining(26), 14 µm cryosections of back skin were fixed with 4% paraformaldehyde/PBS (10 min at room temperature), permeabilised with 0.1% Triton X-100/PBS (10 min at room temperature), blocked with 5% BSA/PBS (1 h at room temperature) and stained with the indicated primary antibodies and 5 µM B-CHP (BIO300, 3Helix) overnight at 4°C. According to the manufacturer’s instructions, the B-CHP probe was heated for 5 min at 80°C before adding it to the primary antibody mixture, which was immediately applied to the tissue sections. Sections were washed four times with PBS and incubated with appropriate secondary antibody and streptavidin–AlexaFluor647 (S32357, Thermo Fisher) for 1 h at room temperature. After additional four washes and DAPI staining slides were mounted as described above.

Confocal microscopy was performed with a Nikon A1 confocal microscope using a 20× objective and brightfield images of H&E and Herovici staining were acquired using a Hamamatsu digital slide scanner with a 40x objective. Image processing was performed with Nikon ND2 Viewer Software, ImageJ (Fiji), Photoshop CS8 (Adobe) and Icy (version 2.1.0.1) software.

### In vivo live imaging and analysis

In vivo live imaging of dermal fibroblasts was performed after 1 and 4 days of acUVB exposure (see experimental design Fig. 4a). Briefly, prior to skin imaging hair follicles were removed with depilation cream (Veet hair removal cream for dry skin) which was applied to the sham and UVR exposed skin area of the lower back and massaged into the skin for approximately 2 min. The area was then washed thoroughly with water, removing hair and cream from the imaging site. Throughout imaging, mice were anesthetised by inhalation of vaporised 1.5% isoflurane (Cp-Pharma) and placed in the prone position in a chamber with body temperature maintained at 37°C via a homeothermic monitoring system (Harvard Apparatus). Additionally, oxygen levels were monitored with the MouseOx Plus (Starr Life Sciences Corp) throughout the imaging sessions using an adult mouse pinch attached to the thigh. Oxygen saturation remained at approximately 99%.

The back skin was stabilised between a cover glass and a thermal conductive soft silicon sheet as previously described(61). Two-photon excitation microscopy was performed with a Zeiss LSM 7MP upright microscope, equipped with a W Plan-APOCHROMAT 20x/1.0 water-immersion objective lens (Zeiss) and a Ti:Sapphire laser (0.95 W at 900 nm; Coherent Chameleon II laser). The laser power used for observation was 2–10%. Scan speed was 4 ls/pixel. The nuclear GFP expression of dermal fibroblasts can be readily detected in PDGFRαH2BEGFP transgenic mice and the autofluorescence of fibrillar collagen can be visualised by the second harmonic signal (SHG) using an excitation wavelength of 770 nm. For time-lapse images z-stacks were acquired every 10 min with a view field of 0.257 mm^2^ in 5 µm steps. A total of 3-6 mice per time point were examined, and the duration of time-lapse imaging was 70-90 min per mouse. Optimisation of image acquisition was performed to avoid fluorescence bleaching and tissue damage and to obtain the best spatiotemporal resolution. Acquired images were analysed with Fiji imaging software (ImageJ, NIH) and Imaris (BitPlane).

Briefly, raw image files (czi) were imported into Fiji where they were subjected to the Correct 3D Drift plugin using channel 1 (collagen) for registration and selecting for sub pixel drift correction(62). The sample drift correction was then manually checked via orthogonal view whereby 3 hair follicles in each sample would be selected in the xz and yz planes and their relative positions measured at time 0 and 80 minutes and an average taken. The final 3D drift corrected time lapse movies were then inputted into Imaris (BitPlane). Within Imaris tracking spots over time was selected using channel 2 (GFP) to follow fibroblast movement over time. Only fibroblasts with a signal quality above 80%, diameter above 8 µm, gap size 3, max distance 15 µm were selected using the autoregressive motion algorithm. This created an animation with spots corresponding to fibroblasts. All spots which corresponded to epidermal noise or hair follicle signal were removed manually for each image. Statistics including cell displacement, position and mean velocity were exported into Excel for further analysis. For calculation of the cell displacement in x-, y-and z-direction, the position of each cell at the start and imaging endpoint was compared and percentage of cells displaced ≥ 5 µm (z-stack imaging step size) was quantified.

### Quantitation and statistical analysis

Statistical analysis was performed with GraphPad Prism 9 software. Unless stated otherwise, data are means ± standard deviation (SD) and statistical significance was determined by unpaired t-test, ordinary One-way or Two-way ANOVA for biological effects with an assumed normal distribution. For unbiased cell identification with DAPI, Ki67, YAP, yH2AX, cCasp3, TOPGFP or PDGFRαH2BEGFP labelling, nuclear staining was quantified using the Spot detector plugin of Icy software (version 2.1.0.1). Similarly, cells labelled with tdTomato in the lineage tracing experiments or stained with Ly6G (neutrophils) were counted per area with the Spot detector plugin of Icy software. To quantify CD45 and CHP staining in tissue sections, mean fluorescence was determined with Icy software and normalised to background. The cell elongation was determined by measuring the maximal tdTomato+ cytoplasm diameter of individual cells in the papillary dermis of Lrig1CreER x Rosa26-tdTomato lineage traced transgenics. Figures were prepared with Adobe Photoshop and Adobe Illustrator (CC2019).

## Acknowledgements

We thank the Nikon Imaging Centre, Dylan Herzog from the Microscopy Innovation Centre here and BSU facility at KCL for expert assistance. Further we would like to thank Matteo Battilocchi (KCL) and Dr Monica Sen (KCL) for assistance with in vivo experiments and the lymph node isolation, respectively. The authors would like to thank Prof Edel O’Toole (Queen Mary University of London) for critically reading the manuscript.

## Author contributions

E.R. conceptualised the study, designed the experiments, performed experiments and analysed the results. G.G. performed in vivo experiments, live imaging and imaging analysis. K.I.K. and K.H.S. performed experiments and analysed data. T.H. assisted with in vivo live imaging. P.L., V.T., I.C. and N.A. performed and analysed the immune cell flow cytometry data. N.J.H., S.M.P. and L.E.R. performed and analysed the human UVR study. E.R. wrote the manuscript and produced the figures. F.M.W. oversaw the study and co-wrote the manuscript. All co-authors commented on the manuscript.

